# Dam Site and Vegetation Selection by Reintroduced Beaver (*Castor canadensis*) on the San Pedro Riparian National Conservation Area, Arizona

**DOI:** 10.1101/2022.06.22.497031

**Authors:** Marcia F. Radke

## Abstract

Beaver (*Castor canadensis*) were reintroduced on the San Pedro Riparian National Conservation Area (SPRNCA) after having been extirpated. The two most obvious effects from this keystone species, dam construction and herbivory, have not been studied on the SPRNCA. The purpose of this research was to determine any preference for dam sites, vegetation preferences, or if beaver may travel longer distances for preferred food or dam materials. Individual dams remained or were rebuilt in the same location on average every 2.4 years. Dam building was disproportionate to availability of sites, with beaver more commonly building dams at the confluence of the San Pedro River and tributaries. The size of areas with impacts to vegetation from beaver herbivory averaged 0.34 ha (2.5 ac), containing an average of 40.7 trees. Beaver exhibited a preference for smaller-sized cottonwood (*Populus fremontii*) trees over larger cottonwood trees and smaller or larger Goodding’s willow (*Salix gooddingii*), both in terms of whether trees were eaten and left standing or felled, and distance beaver travelled from water to the tree. Management implications include continued research on effects of beaver, management of invasive plant species, use of cottonwood genotypes with high tannin levels during restoration projects, further augmentation of beaver, use of beaver-dam analogues, and continued closure to hunting and trapping of beaver.

## INTRODUCTION

Beaver (*Castor canadensis*) were reintroduced on the San Pedro Riparian National Conservation Area (SPRNCA), after having been extirpated by fur trappers by 1894 (Bailey 1971). Beaver dams may increase storage capacity and lead to greater flows during dryer periods (Parker 1986), elevate water tables through groundwater recharge (Johnston and Naiman 1987), and increase hyporheic flows (Kasahara and Wondzell 2003; Westbrook et al. 2006). While beaver dams may improve water storage, beaver herbivory may have a significant impact on vegetation where foraging occurs and dam material is collected. I hypothesize that beaver may show preferences for dam sites, different species of vegetation at dam sites, and that beaver may travel longer distances for preferred food or dam materials.

## STUDY AREA

The SPRNCA, located in southeastern Arizona approximately 85 km southeast of Tucson, was established in 1988 by the U.S. Bureau of Land Management. The SPRNCA encompasses 56,431 acres and contains 51 miles of the San Pedro River immediately north of Sonora, Mexico. Fremont cottonwood /Goodding’s willow gallery forest occurs over much of the river’s length, with increasing cover of the exotic tamarisk (*Tamarix ramosissima*) in the northern ephemeral section.

## METHODS

Data analyzed for beaver dams included years from 2000-2010. Beaver dams were considered active for the hydrologic year after monsoon (October) up to monsoon the following calendar year (June); dams were usually removed through monsoonal flooding events in July-September. ArcGIS was used to map site fidelity and expansion of dams. In order to determine preferential selection of dam sites, the number of dams at tributaries was compared to the number of dams not at tributaries. Due to GPS accuracy, dam site selection was considered to be at the same site if dams occurred within 50 m (164 ft) of prior years’ locations. An approximate 51 km (31.7 mi) of the SPRNCA was analyzed for a total of 1,020 possible dam sites. A chisquare goodness-of-fit test using Yates correction for continuity (Zar 1999) was used to determine if dam sites were used in proportion to availability. Confidence intervals were constructed around the proportion of observed use to determine any selection.

Herbivory sites were identified during 2008 to 2009 as all areas where beavers foraged on land from a dam site. Once an active site was located, data was collected on all vegetation species that showed sign of beaver herbivory at indefinite distance from the dam. Herbivory sign included everything from single incisor marks to entire trees that were felled. Site size was determined using the UTM coordinates of each plant with herbivory, and a polygon was created on ArcGIS to determine the area. Trees with herbivory were characterized as standing or felled, and measurements of diameter at breast height (DBH), height, and species were obtained. Vegetation species with beaver herbivory included Fremont cottonwood, Goodding’s willow, or seep willow (*Baccharis salicifolia*). DBH and distance to water was measured using a tape, and height of tree using a clinometer. The DBH of multiple stems or trunks from the same rooted plant was obtained by adding multiple trunks or stems together. Size class I included Fremont cottonwood and Goodding’s willow with ≤ 48 cm (18.9 in) DBH, while size class II included Fremont cottonwood and Goodding’s willow with >48 cm (18.9 in) DBH.

The *a priori* significance threshold was 0.05, and design for analysis of DBH and distance to water for standing and felled trees was a parametric three-factor analysis of variance (ANOVA). After data collection, I tested for normality for n≥20 using the D’Agostino-Pearson K^2^ test for normality (Zar 1999). Assumptions for parametric ANOVA were not met if a significant difference from normality existed for each data set. In this case, I used a nonparametric Kruskal-Wallis ANOVA with tied ranks, and any significant difference between groups was determined using nonparametric Mann-Whitney pairwise comparisons.

## RESULTS

### Dam site expansion and fidelity

The total number of beaver dams steadily increased from five in 2000 to 39 during 2010, with a cumulative total of 235 dams in 170 locations, and an average of 21.4 dams per year. Dam site fidelity ranged from one to eight years including both consecutive and non-consecutive years. Of 235 dams documented, 166 locations (71%) were used for one year, 19 dam locations (34%) were used for two years, 15 (27%) were used for three years, eight (14%) were used for four years, eight (14%) were used for five years, one (2%) was used for six years, three (5%) were used for seven years, and 2 (4%) were used for eight years (Table 1). The average number of years individual dams remained in the same location after monsoon season or were rebuilt at the same location was 2.4.

**Table 1.** Number of years in same location that beaver dams have remained active, 2000 to 2010, San Pedro Riparian National Conservation Area.

| Number of Years | Number of Dams (%) |
| --- | --- |
| 1 | 36 |
| 2 | 19 (34) |
| 3 | 15 (27) |
| 4 | 8 (14) |
| 5 | 8 (14) |
| 6 | 1 (2) |
| 7 | 3 (5) |
| 8 | 2 (4) |

### Dam site selection

Analysis of the number of beaver dams at tributary washes versus the number of dams not at washes indicated that the samples were homogenous (χ^2^=16.53, df=10, 0.05<P<0.10). The pooled data indicated there was a significantly higher number of dams constructed at tributary washes versus dams not constructed at washes (Table 2; χ^2^=1148.84, df=1, P<0.001).

**Table 2.**
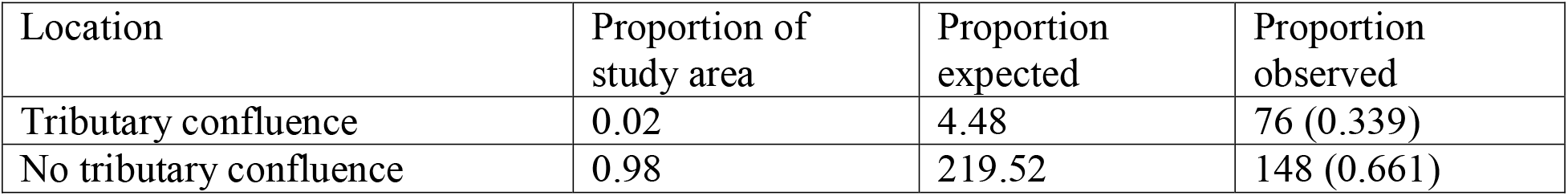
Selection of tributary confluences by beaver for dam sites, San Pedro Riparian National Conservation Area, 2000-2010. Chi-square value = 1148.84; significant at P<0.001.

### Dam site vegetation selection

The size of areas with impacts to vegetation from beaver herbivory (n=6) ranged from 0.06 – 0.81 ha (0.15 – 2.0 ac) (mean=0.34 ha or 2.5 ac), containing from 11 to 134 trees (mean=40.7). In each area, the number of Freemont cottonwood with herbivory ranged from 1 to 70, the number of Goodding’s willow with herbivory ranged from 0 to 43, and the number of seep willow with herbivory ranged from 0 to 21. Velvet mesquite (*Prosopis velutina*), hackberry (*Celtis reticulata*), and tamarisk were documented with beaver herbivory one time for each species.

### Vegetation selection by species and size

The size of standing single trunk Freemont cottonwood with beaver herbivory ranged from 10.8 to 101.9 cm (4.3 to 40.1 in) DBH, and from 6.4 to 29.8 m (21.0 to 97.8 ft) in height. The size of single trunk Goodding’s willow with beaver herbivory ranged from 4.7 to 101.5 cm (4.9 to 40.0 in) DBH, and from 1.0 to 21.8 m (3.3 to 71.6 ft) in height. Seep willow ranged from 1.9 cm (0.7 in) (single stem) to 31.8 cm (12.6 in) DBH, and from 0.8 to 3.3 m (2.6 to 10.8 ft) in height.

The size of felled single trunk Freemont cottonwood ranged from 9.9 to 41.4 cm (3.9 to 16.3in) DBH, and from 4.8 to 31.5 m (15.7 to 103.3 ft) in height. The size of single trunk Goodding’s willow ranged from 1.2 to 25.5 cm (0.5 to 10.0 in) DBH, and from 1.0 to 15.0 m (3.3 to 49.2 ft) in height. Seep willow ranged from 1.6 (0.6 in) (single stem) to 17.5 cm (6.9 in) DBH, and from 1.5 to 3.3 m (4.9 to 10.8 ft) in height.

For all areas with beaver herbivory of standing and felled trees, the mean DBH and height for Freemont cottonwood, Goodding’s willow, and seep willow were 31.8 cm (12.6 in) and 18.4 m (60.4 ft), 21.2 cm (8.3 in) and 8.3 m (27.2 ft), and 7.8 cm (3.0 in) (including all stems from the same rooted plant) and 2.1 m (6.9 ft), respectively.

The analysis of DBH and standing or felled Fremont cottonwood and Goodding’s willow showed significant differences (Kruskal-Wallis test, Hc=72.76, df=3, P=2.587E^-15^), with five significant comparisons between the two species of trees and whether they were standing or felled. Higher numbers of standing trees were directed at larger size (Figure 1; Mann-Whitney pairwise comparisons, Bonferroni corrected).

**Figure 1.**
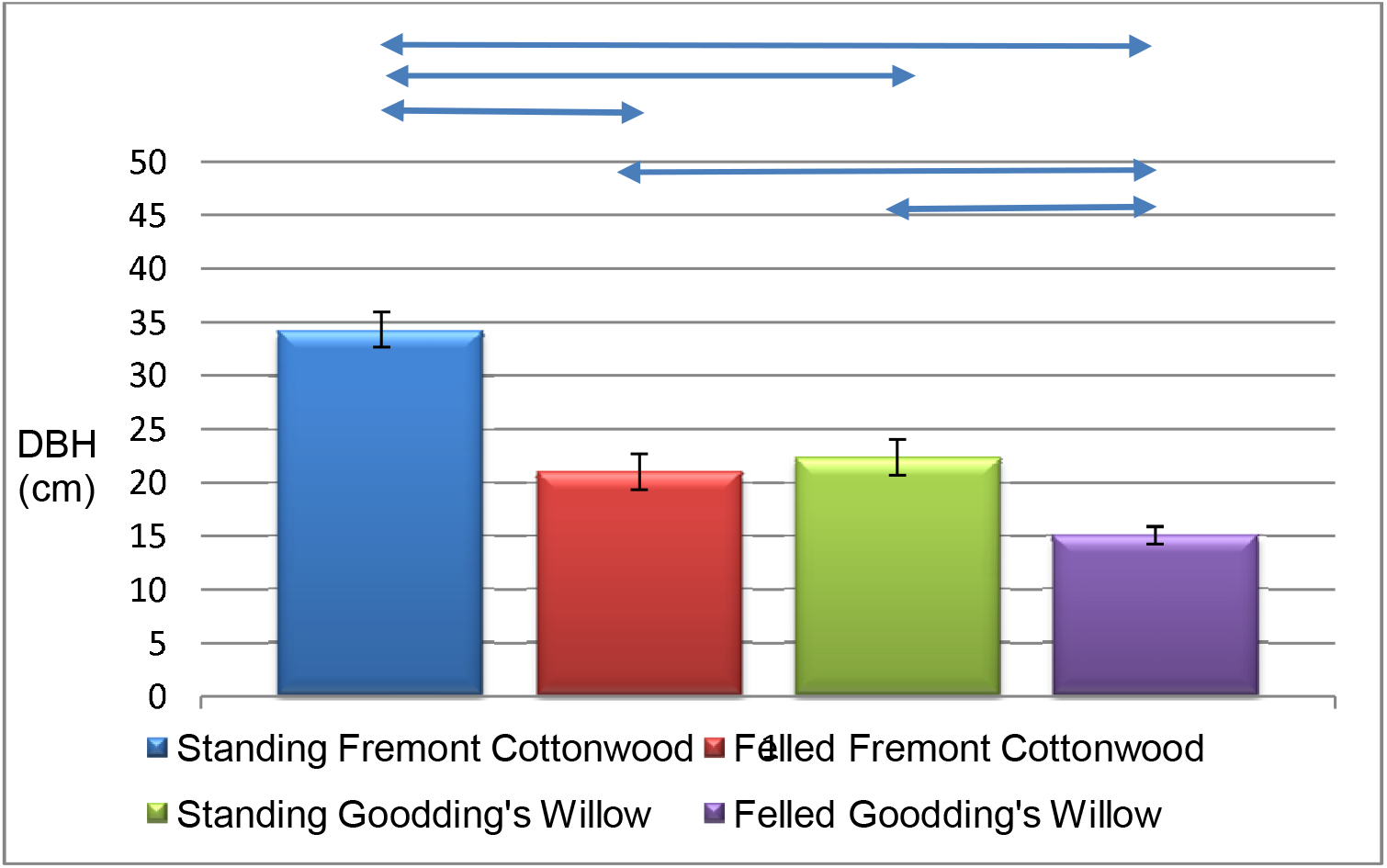
Mean (± standard error) diameter at breast height (cm; DBH) Fremont cottonwood and Goodding’s willow left standing or felled by beaver, San Pedro Riparian National Conservation Area, 2008-2009. Significance is indicated by arrow between species and size class.

The mean DBH of standing Fremont cottonwood was significantly larger than the mean DBH of felled Fremont cottonwood (P=0.0001), the mean DBH of standing Fremont cottonwood was significantly larger than the mean DBH of standing (P=3.224E^-06^) or felled Goodding’s willow (P=1.204E^-12^), the mean DBH of felled Fremont cottonwood was significantly larger than the mean DBH of felled Goodding’s willow (P=0.0356), and the mean DBH of standing Goodding’s willow was significantly larger than the DBH of felled Goodding’s willow (P=0.00457). No significant difference existed between the comparison between the mean DBH of felled Fremont cottonwood and standing Goodding’s willow (P=1.0).

### Vegetation selection by distance to water

Mean distance from the San Pedro River to vegetation with herbivory was 5.9 m (19.4 ft) for Freemont cottonwood, 3.8 m (12.5 ft) for Goodding’s willow, 2.1 m (6.9 ft) for seep willow, and 4.7 m (15.4 ft) for all species combined. Mean distance from the San Pedro River to Fremont cottonwood or Goodding’s willow with beaver herbivory was significantly different (Figure 2; Kruskall-Wallis test, F=8.457, df=217, P<0.01). Significant differences existed between four comparisons between distance to water between Fremont cottonwood size class I and size class II (Mann-Whitney pairwise comparison, P=0.043), between Fremont cottonwood size class I and Goodding’s willow size class I (P=0.0013), between Fremont cottonwood size class I and Goodding’s willow size class II (P= 3.154E^-05)^, and between Fremont cottonwood size class II and Goodding’s willow size class II (P=0.0005). The mean distance was not significantly different between cottonwood size class II and Goodding’s willow size class 1 (Mann-Whitney pairwise comparison, P=0.052), or between Goodding’s willow size class I and Goodding’s willow size class II (Mann-Whitney pairwise comparison, P=0.066).

**Figure 2.**
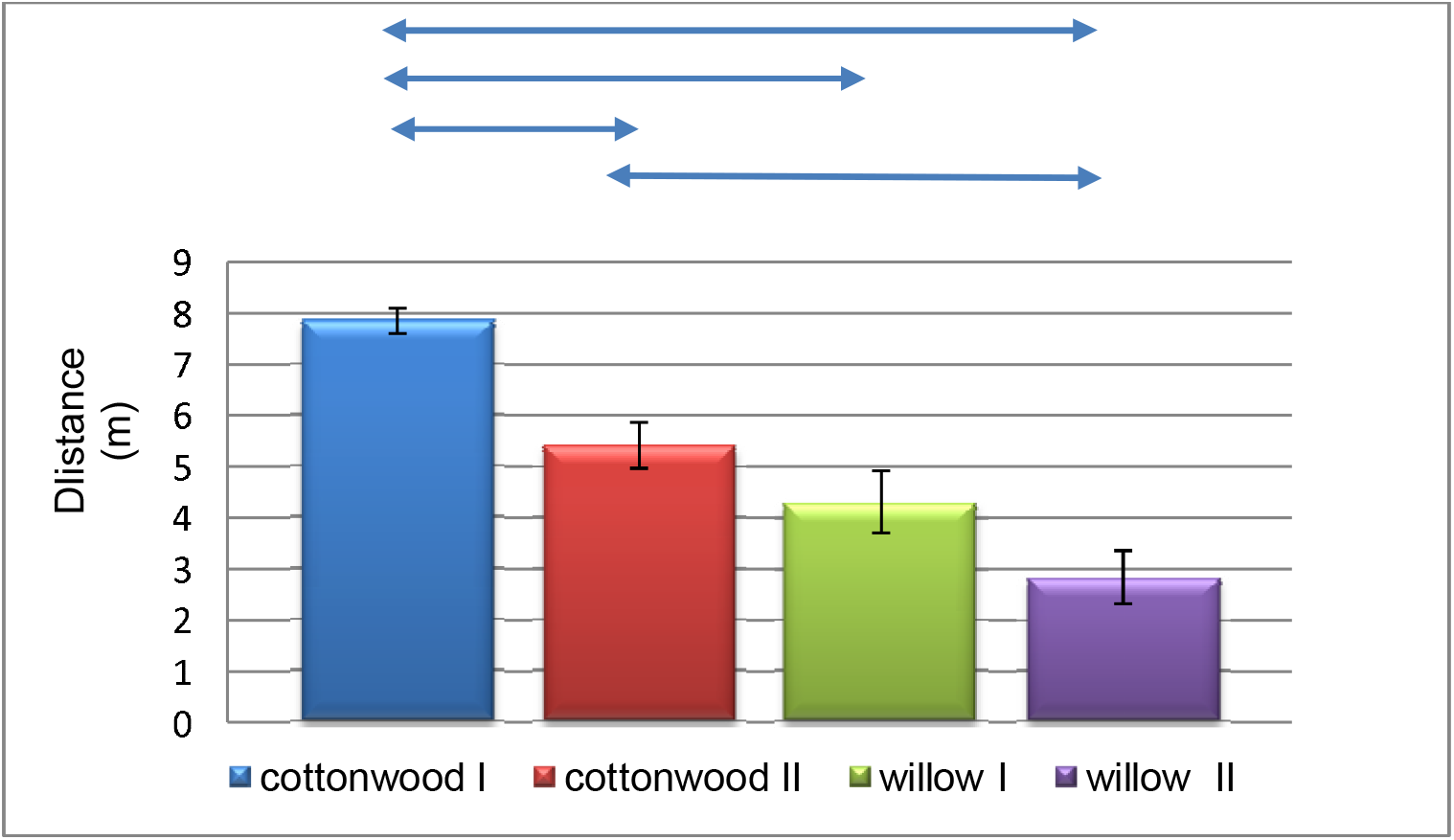
Mean (± standard error) distance (m) from water to size class I and size class II Fremont cottonwood and Goodding’s willow with beaver herbivory, San Pedro Riparian National Conservation Area, 2008-2009. Significance is indicated by arrow between species and size class.

## DISCUSSION

### Dam site expansion and fidelity

Initial accounts of beaver and dams after the initial reintroduction were within the San Pedro River’s main stem. The number of dams on SPRNCA grew quickly, with hundreds of dams documented in approximately 15 miles of perennial water. A few dams were not consistently used at perennial Lewis Springs (2010), or in intermittent reaches of the river. The Upper San Pedro River watershed, at Represo Los Fresnos, Sonora, Mexico, had beaver accounts quickly following the initial reintroduction. Beaver herbivory was noted in Bear Creek on the south side of the Huachuca Mountains during 2010 and 2011 and, for the Upper Santa Cruz watershed, beaver sign was noted during 2012 near San Lazaro, Sonora, Mexico (Trevor Hare, personal communication, December 18, 2017). Beaver sign was noted in the San Rafael Valley during 2016 (Doug Duncan, U.S. Fish and Wildlife Service, personal communication, December 18, 2017). During 2014, a beaver was observed and photographed in a stock tank on Campini Mesa, between the San Pedro and Santa Cruz watersheds (Traci Swift, personal communication, August 5, 2014), where overland travel by beaver would have been necessary. During 2017, beaver dams were noted in the Babocomari River, a perennial tributary to the San Pedro River (Laura Norman, U.S. Geological Survey, personal communication, April 13, 2017).

A review of historical references noted several accounts of beaver in the Santa Cruz watershed, including the Rillito, Cienega, and Sonoita tributaries (Fish and Gillespie 1987). Historic climate and habitat conditions were likely more conducive to beaver than current ongoing drought conditions in southeast Arizona. With beaver emigrating to the Santa Cruz watershed from the San Pedro watershed under current conditions, it appears beaver must have also been able to do so historically.

Channel entrenchment occurred historically along the San Pedro River, resulting in large amounts of water with high velocity funneling through the channel during large flood events. Small or isolated flood events may not wash out dams, but larger flood events wash out many, if not all, beaver dams. For example, heavy monsoonal rains during 2008 removed all 33 beaver dams built during 2007-2008, and the 2010 monsoon season again removed all 39 beaver dams built during 2009 and 2010. Thus, beaver dam site fidelity on SPRNCA currently appears to be mainly affected by rain events which trigger high flood flows in the river.

Beaver behavior may also be affected by monsoonal flooding. Beaver leave their parent colony at approximately two years of age (Collen and Gibson (2001). Subadult beaver may use these flood events to leave their natal colony and disperse, both upstream and downstream, in order to find their own territory and mate.

### Dam site selection

Building of dams by beaver was disproportionate to availability of sites, with beaver more often building dams at the confluence of the San Pedro River and tributaries. Tributaries create rock, gravel, and sand bars, resulting in natural ponding of water in the river channel upstream of the tributary, with or without beaver dams. In some locations, tributaries prevent downstream surface water flow, but also contribute significant surface and/or groundwater which serves to slow water by creating a backwater effect. Shallower depths and shorter perpendicular distances of surface water over the tributary bars may be present, resulting in beaver dams that are shorter in height and length with less material needed for dams. If a dam is needed at all, beaver may expend less energy at tributary intersections in tree-felling, moving dam materials, and dam construction, with more energy available for other activities, such as feeding, mating, and raising young. Similarly, Rosell et al. (2005) noted that size of dams constructed by beaver may be largely dictated by topography of the site and availability of building materials. Additionally, selection of colony sites near stream bifurcations may increase the shoreline available for foraging near the colony (Boyce 1981). New pond creation becomes limited to less desirable sites (Johnston and Naiman 1990), so that preferential use of tributary areas by beaver may change over time.

### Vegetation selection

For the purposes of this discussion, size class I (≤ 48cm) is denoted as “smaller” and size class II (>48cm) is denoted as “larger” sized trees. A partiality by beaver was exhibited for smaller-sized cottonwood or willow compared to larger-sized trees of the same species, and a partiality for cottonwood even over smaller willow. Analyses of distance to water beaver travelled for different size classes of cottonwood and willow revealed similar results. Beaver travelled significantly longer distances to reach smaller, compared to larger-sized cottonwood, indicating a willingness by beaver to expend more energy and time travelling to smaller-sized cottonwood. Similarly, beaver travelled significantly longer distances for smaller cottonwood, rather than smaller and larger-sized willow, demonstrating a preference for cottonwood even when less energy and time could have been spent on travelling to both small and larger-sized willow. A significant difference existed between larger cottonwood and larger willow, with beaver travelling further distances for larger cottonwood than willow, indicative of a partiality for even large cottonwood over willow.

Reasons for beaver preference between cottonwood and willow may involve several factors. Riparian zones form distinct vegetative mosaics with contrasts in structure related to the underlying physical environment (Balian and Naiman 2005). Within the SPRNCA, the mean annual maximum depth to groundwater is approximately 2 m (6.6 ft), with willow occurring at a slightly shallower maximum depth (Leenhouts et al. 2006). Due to the requirements for higher ground water levels, willow generally occur closer to the water’s edge, while cottonwood may occur further from the surface water where they established during a different flow regime. Seep willow generally occurs close to the edge of the San Pedro River where it is readily available, however, it does not provide large woody materials needed for dam building. This may explain why many areas of beaver herbivory contained seep willow without herbivory where it occurred concurrently with cottonwood and willow. The willingness of beaver to travel farther from the water for cottonwood, even with availability of closer willow, indicates there may be a texture, taste, and/or nutritional preference for smaller cottonwood that compensates for the higher energetic costs in travel.

Several chemical groups may influence herbivory, and concentrations may vary between large and small cottonwood trees (Meyer and Montgomery 1987). Beaver may show a predilection for smaller cottonwood based on lower concentration of defensive chemicals that vary with genetics (Woolbright et al. 2008). It is also possible that smaller cottonwood trees provide better textured or lighter weight materials for dam construction. Cottonwood is approximately 100 times lighter in weight, for both fresh green and air-dried seasoned wood, than willow species. Felling and handling smaller cottonwood trees may expend less energy. Reduced energetic demands and perhaps better nutritional values from processing smaller cottonwood may result in enhanced capability to protect territories, mate, and produce young.

Fremont cottonwood will resprout from the stump (McGinley and Whitham 1985). In this study, cottonwood resprouts are now approximately 3 m tall. Recruitment of cottonwood and willow is not only occurring from resprouts caused by beaver herbivory, however. Many young cottonwood and willow trees, approximately three to five years of age, are present in all river reaches. These young trees appear to be growing on seed beds created by large, scouring floods that occurred during 2013 and 2014. Therefore, recruitment of the native gallery forest continues, despite beaver herbivory, with scouring flows responsible for recruitment.

## MANAGEMENT IMPLICATIONS FOR THE SPRNCA

1. Monitoring of vegetation and other wildlife should continue to address effects of beaver reintroduction.
2. Because longer-term browsing by beaver may cause replacement by non-preferred species (Donkor and Fryxell 2000), selective vegetation control of tamarisk should continue.
3. Cottonwoods high in tannin and unpalatable to beaver may be optimal choices for restoration plantings (Durben and Walker 2010).
4. Augmentation of the beaver population should be considered if insufficient genetic diversity (i.e. founder effect) is suspected.
5. Beaver dam analogues (BDAs) have been used in other locations as a riparian restoration technique (Pollack et al. 2014), which should be considered if natural beaver dams do not develop at densities sufficient to cause aggradation of the entrenched channel.
6. In order to allow for future increase in the beaver population, continued closure to hunting and trapping of beaver should occur.

